# Seasonal Light and Temperature Timing in a Stoichiometric Food Web

**DOI:** 10.64898/2026.09.13.751257

**Authors:** Ramiro Ramirez

**Affiliations:** University of Texas at San Antonio Health Science Center

**Keywords:** ecological stoichiometry, phenological mismatch, periodic environment, Floquet multiplier, benthic–pelagic coupling, *Daphnia*, food quality

## Abstract

Seasonal food-web interactions can depend on whether consumer performance is high when food quantity and elemental quality are favorable. We extend a closed-phosphorus model containing pelagic and benthic producers, variable producer phosphorus quotas, and a shared *Daphnia* grazer by allowing annual light and temperature cycles to have an adjustable phase difference. The previous light-seasonality preprint has a nonisolated grazer-free boundary. We parameterize its annual periodic extension by the fraction of producer phosphorus in phytoplankton and derive the unique positive annual producer orbit for every fixed allocation. Linearization in the rare-grazer direction then gives an exact conditional Floquet exponent. Its decomposition into a mean-rate term and a covariance term identifies the effect of seasonal timing. A reconstructed descriptive thermal proxy uses quasi-acclimated filtration-capacity means from the official Müller et al. dataset; it supplies only a relative response shape over 15–25 °C, not an absolute ingestion calibration. In a representative configuration, changing phase while preserving the annual light and temperature distributions changes the invasion exponent from −0.00382 to 0.01486 day^−1^, with annual multipliers 0.248 and 227, respectively. The constant-mean-ingestion exponent is positive, so the negative case is generated by adverse timing covariance. Sign reversal persists across a range of phosphorus allocations, but not across the entire boundary family. The result is a local invasion criterion for specified grazer-free cycles, not a theorem of global persistence or extinction. Within this parameterized boundary problem, relative seasonal timing can change the sign of infinitesimal consumer growth.

## 1 Introduction

Light and phosphorus jointly determine both the amount and elemental quality of producer biomass in aquatic food webs. Under high light relative to phosphorus, producers can accumulate carbon faster than phosphorus, reducing their value to phosphorus-demanding consumers [2]. Stoichiometric producer–grazer models therefore distinguish food quantity from food quality and can produce consumer responses that are not predicted by producer biomass alone [1, 3].

Seasonal timing adds another dimension to this problem. Light changes producer carrying conditions, whereas temperature can change consumer feeding and other physiological rates. Periodic stoichiometric studies show that seasonal light can change population dynamics and generate periodic or quasiperiodic behavior [3, 5], while general periodic stoichiometric threshold results are also available [4]. Temperature and nutrient supply can interact to redistribute biomass or destabilize consumer–resource systems [6, 7, 13]. Experiments and modeling with *Daphnia* further show that resource storage, forcing frequency, covariance, resource-peak phase, and seasonal food quality can change consumer performance [8, 9, 10]. Food quality can also change the magnitude and direction of *Daphnia* growth responses to thermal fluctuations [12]. Recent individual-level work crosses temperature variation with food quantity and quality when analyzing growth and maturation [11]; it does not study phase-shifted population invasion. Climate change can therefore create nutritional phenological mismatches [14]. A phase shift can matter without changing annual mean environmental conditions.

Rare-consumer invasion calculations are established in seasonal food-web theory [15], while the frequency of periodic forcing can independently change consumer–resource stability [16]. In a different deterministic and stochastic resource framework, desynchronizing substitutable seasonal resources can reduce extinction risk [17]. Our goal is not to claim that phase effects or periodic invasion analysis are new in themselves. Instead, we ask how they operate in a specific stoichiometric food web that couples phytoplankton and periphyton through a shared grazer and habitat-specific phosphorus access.

The underlying six-state model was developed in the author’s dissertation [18] and subsequently analyzed under constant and seasonally varying light [19]. Dissertation Chapter 4 explored temperature-dependent ingestion under constant light, and its conclusion proposed combined light–temperature forcing with preliminary illustrations. Here we develop that proposed extension through a new annual phase experiment, a reconstructed thermal proxy, and an analytical criterion for growth from rarity.

The paper has three objectives. First, we extend the preprint’s known continuum of grazer-free equilibria to annual resident cycles indexed by producer phosphorus allocation. Second, we derive the positive periodic producer orbit and the conditional rare-grazer multiplier on every member of that family. Third, we use the resulting exponent to test whether changing only the phase between annual light and temperature can reverse rare-grazer growth. The central result is a conditional sign reversal and an exact covariance explanation. It does not establish global persistence, attribute the effect uniquely to separate nutrient pools, or provide a calibrated forecast for a natural lake.

## 2 Seasonally forced stoichiometric model

Let *x* and *w* denote phytoplankton and periphyton carbon biomass, *y* the grazer biomass, *Q*_*x*_ and *Q*_*w*_ the producer phosphorus-to-carbon ratios, and *P*_*B*_ and *P*_*p*_ the benthic and pelagic dissolved-phosphorus pools. The grazer has fixed phosphorus-to-carbon ratio *θ*_*y*_. Total phosphorus is conserved:

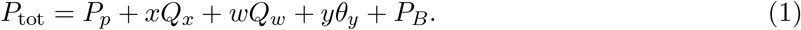

1) Producer phosphate uptake and specific growth are

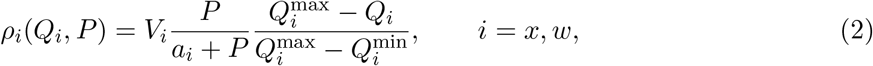

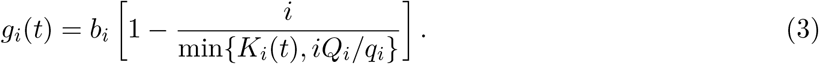

The grazer conversion factor is

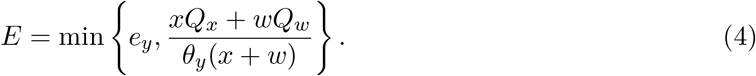

The six-state quota formulation is

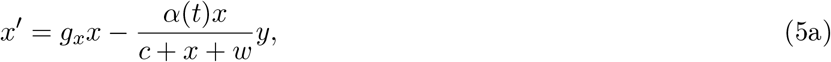

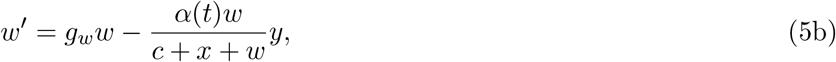

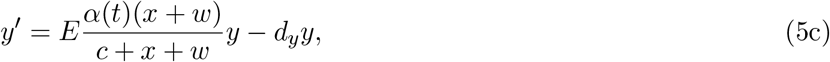

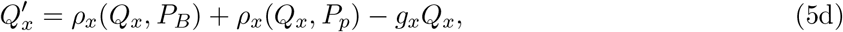

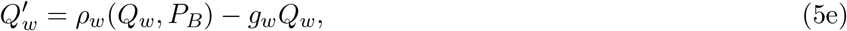

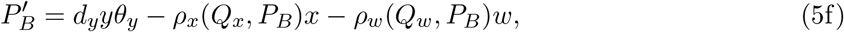

with *P*_*p*_ recovered from Eq. (1). Phytoplankton accesses both dissolved pools, whereas periphyton accesses only the benthic pool. These pathways are model assumptions. The construction, phosphorus accounting, boundedness conditions, equilibria, and constant-light bifurcation structure are given in [19]; here we focus on the new periodic boundary problem.

We assume positive model parameters, 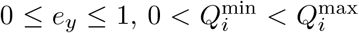, and 0 *≤ ε <* 1. We use common annual light forcing

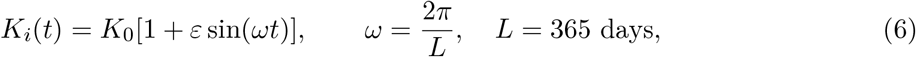

where 0 *≤ ε <* 1. Temperature has the same period and an adjustable phase:

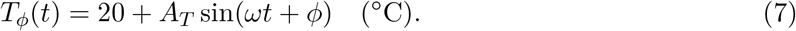

Thus *ϕ* = 0 aligns the temperature and carrying-capacity peaks, and positive *ϕ* advances the temperature peak relative to light. Unlike the 182.5-day light-only numerical reconstruction in [19], Eq. (6) represents one annual cycle. This is a new modeling choice, not a reinterpretation of the submitted calculation.

### 2.1 Thermal-response proxy

We reconstruct the thermal proxy from source measurements rather than reuse the dissertation calibration. That calibration used an invalid temperature transformation and observations measured after return to 20 °C rather than at the listed ambient temperatures. The surviving combined-forcing script also retains constant ingestion, so the preliminary combined-forcing figures are not used here. The accompanying provenance notes document these discrepancies.

Müller et al. measured *Daphnia magna* filtration capacity under temperature exposure and reversal treatments [20]. Their official dataset gives quasi-acclimated mean filtration capacities and reported standard deviations of 1.317 *±* 0.035, 2.345 *±* 0.065, 2.9651 *±* 0.057, and 1.888 *±* 0.040 mL individual^−1^ hour^−1^ at 11, 15, 20, and 25 °C, respectively [21]. The 29 °C exposure was lethal and does not supply a quasi-acclimated value. We fit the descriptive curve

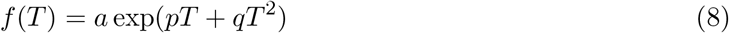

by weighted nonlinear least squares, treating the reported treatment standard deviations as observation-scale weights. We transfer only its relative shape to the food-web model:

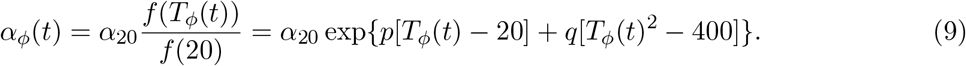

Filtration capacity is not identical to the maximal ingestion coefficient in Eq. (5). Accordingly, *α*_20_ = 0.8 day^−1^ retains the existing model scale, while *f*(*T*)*/f* (20) is an exploratory quasi-acclimated filtration proxy. Four aggregated points and three fitted parameters provide little freedom for model checking; the fitted uncertainty should not be interpreted as population-level physiological uncertainty.

## 3 Grazer-free periodic family and conditional invasion

It is useful to write producer phosphorus contents as *P*_*x*_ = *xQ*_*x*_ and *P*_*w*_ = *wQ*_*w*_. Consider the grazer-free invariant face

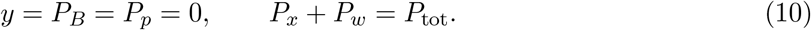

For a fixed allocation *r ∈* (0, 1), set

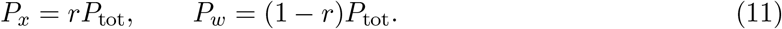

Both phosphorus contents remain constant on this face. Define

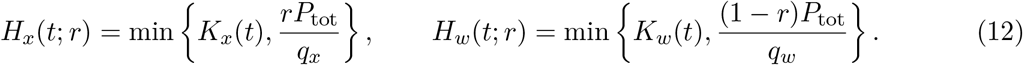

Then each producer satisfies the periodically forced logistic equation

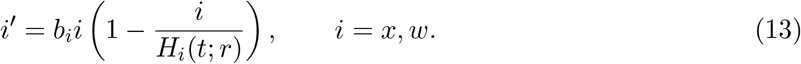

### Theorem 3.1

(Boundary orbit and conditional grazer multiplier). *Under the positive model-parameter assumptions above, fix* 0 *≤ ε <* 1 *and r ∈* (0, 1). *The reduced system in Eq*. (13) *has a unique positive L-periodic producer orbit* (*x*_*r*_(*t*), *w*_*r*_(*t*)) *for that allocation. This orbit belongs to the biological quota state space if its recovered quotas satisfy* 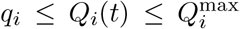. *Sufficient conditions are* 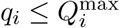*and* 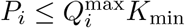, *where K*_min_ = *K*_0_(1 − *ε*) *and P*_*i*_ *is given by Eq*. (11). *Let S*_*r*_ = *x*_*r*_ + *w*_*r*_ *and*

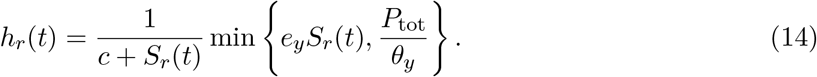

*The grazer variational equation along this orbit is*

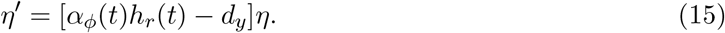

*Its conditional transverse multiplier is M*_*y*_(*r, ϕ*) = exp[*L*Λ(*r, ϕ*)], *where*

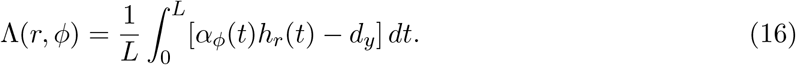

*Thus* Λ *>* 0 *implies net growth and* Λ *<* 0 *net decay over each full period for an infinitesimal grazer along that specified orbit*.

*Proof*. Let *z*_*i*_ = 1*/i*. Equation (13) becomes

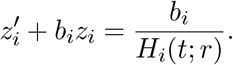

Because *H*_*i*_ is positive and *L*-periodic, the unique periodic solution is

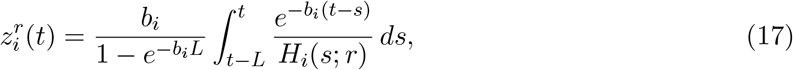

which is positive; 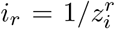 gives the required producer orbit. The difference of two solutions of the linear equation is multiplied by 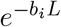 *<* 1 over a period, establishing uniqueness of its periodic solution. The kernel in Eq. (17) has total weight one, so

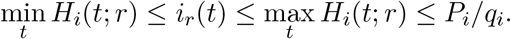

Consequently,

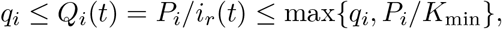

which proves the sufficient quota conditions.

On the face, uptake vanishes and the product rule gives 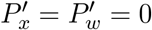. The dissolved pools and *y* remain zero, which justifies the invariant reduction. Along the orbit, *x*_*r*_*Q*_*x*_ + *w*_*r*_*Q*_*w*_ = *P*_tot_. Write the grazer equation as *y*′ = *A*(*t, U*)*y*, with *U* the full state. Near this positive producer orbit, *A* is locally Lipschitz even at the minimum-function switches. Since the resident state *U*_*r*_ has *y* = 0,

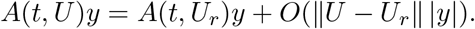

The first-order grazer equation is therefore Eq. (15), without requiring a differentiable full-system Jacobian at every switch. Direct integration of this continuous scalar periodic equation gives the stated multiplier.

The recovered orbit represents an admissible member of the biological quota state space when 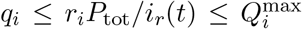 throughout the period, where *r*_*x*_ = *r* and *r*_*w*_ = 1 − *r*. The multiplier is conditional because the grazer-free cycles form a neutral family indexed by *r*. Its sign does not determine stability tangent to that family, nor does it establish uniform persistence or global extinction in the full nonlinear system.

### Corollary 3.2

(Timing covariance). *For fixed r and A*_*T*_, *changing ϕ preserves the annual distribution and mean* 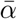 *of α*_*ϕ*_. *The exponent decomposes exactly as*

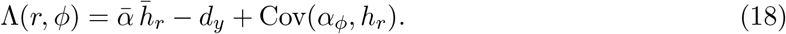

*Here* 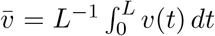 *dt and* Cov 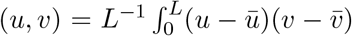 *dt. Consequently, phase changes* Λ *only through the covariance term*.

*Proof*. Changing *ϕ* translates the periodic function *α*_0_ in time, preserving its full-period mean. The resident food factor *h*_*r*_ is independent of *ϕ* because temperature affects only ingestion and the resident grazer is absent. Expanding the centered product in the covariance definition gives 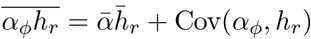, and Eq. (16) yields the identity.

The quantity 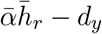 is the constant-mean-ingestion control. It preserves the annual mean of the nonlinear thermal rate. A separate comparison between 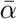 and *α* 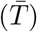 quantifies the nonlinear temperature-to-rate, or Jensen, component.

## 4 Numerical methods

Table 1 lists the parameters inherited from the checked light-seasonality model. We specialize 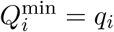 numerically, while retaining their distinct roles in Eqs. (2) and (3). We use *K*_*x*_ = *K*_*w*_ = *K*_0_[1+*ε* sin(*ωt*)]. The primary configuration is *K*_0_ = 1.5 mg C/L, *ε* = 0.7, *A*_*T*_ = 5 °C, and *r* = 0.45. This gives *P*_*x*_ = 0.0135 and *P*_*w*_ = 0.0165 mg P/L. The ecological temperature cycle remains between two observed quasi-acclimated temperatures, 15 and 25 °C, and does not extrapolate beyond that interval.

**Table 1:** Parameters used in the seasonal phase analysis.

| Parameter | Description | Value | Units |
| --- | --- | --- | --- |
| $b_x, b_w$ | Maximal producer growth rates | 1.1, 1.3 | day<sup>-1</sup> |
| $\alpha_{20}$ | Ingestion coefficient at $20^\circ\text{C}$ | 0.8 | day<sup>-1</sup> |
| $c$ | Ingestion half-saturation biomass | 0.25 | mg C/L |
| $e_y$ | Maximal production efficiency | 0.8 | unitless |
| $d_y$ | Grazer loss rate | 0.25 | day<sup>-1</sup> |
| $\theta_y$ | Grazer P:C ratio | 0.03 | mg P/mg C |
| $q_x, q_w$ | Producer minimum growth quotas | 0.0038 | mg P/mg C |
| $Q_x^{\text{min}}, Q_w^{\text{min}}$ | Uptake-normalization quotas | 0.0038 | mg P/mg C |
| $P_{\text{tot}}$ | Total phosphorus | 0.03 | mg P/L |
| $a_x, a_w$ | Phosphate half-saturation concentrations | 0.0015 | mg P/L |
| $V_x, V_w$ | Quota uptake rates | 0.2, 0.3 | mg P/(mg C day) |
| $Q_x^{\text{max}}, Q_w^{\text{max}}$ | Producer upper quotas | 2.5 | mg P/mg C |

We computed the periodic initial value in Eq. (17) by adaptive quadrature and integrated the linear *z*_*i*_ equations with DOP853 at relative tolerance 2 *×* 10^−11^, absolute tolerance 2 *×* 10^−13^, and a two-day maximum step. Annual averages use a 0.25-day grid. The primary amplitude–phase map uses *A*_*T*_ = 0–5 °C in 0.125 °C increments and phase in 2° increments. The reported extrema use a 0.25° phase grid. The allocation map uses *r* = 0.05–0.95 in increments of 0.01. A coarser comparison scans *K*_0_ = 1.4, 1.5, 1.6, 1.7 and *r* in increments of 0.05.

We independently re-integrated the primary producer orbit with DOP853 and LSODA at relative tolerance 10^−10^ and absolute tolerance 10^−12^. We also introduced grazer densities 10^−8^, 10^−10^, and 10^−12^ into the full nutrient-content system, transferring the grazer’s phosphorus from *P*_*x*_ to preserve Eq. (1). These one-year integrations check the linear exponent at both the favorable and unfavorable phase extrema. All phase comparisons use the same annual light time series and the same annual temperature distribution.

## 5 Results

### 5.1 Reconstructed relative thermal response

The weighted fit is

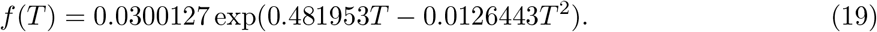

The local weighted-fit covariance gives nominal coefficient standard errors of 0.00506, 0.01947, and 0.000530, respectively. With four aggregated means and three fitted coefficients, these values are not used as inferential or population-level physiological uncertainty. The fitted optimum is 19.06 °C and the descriptive *R*^2^ is 0.9964 (Figure 1). With only one residual degree of freedom, the high *R*^2^ provides little evidence about predictive performance. Equation (19) is used only through the relative proxy in Eq. (9).

**Figure 1:**
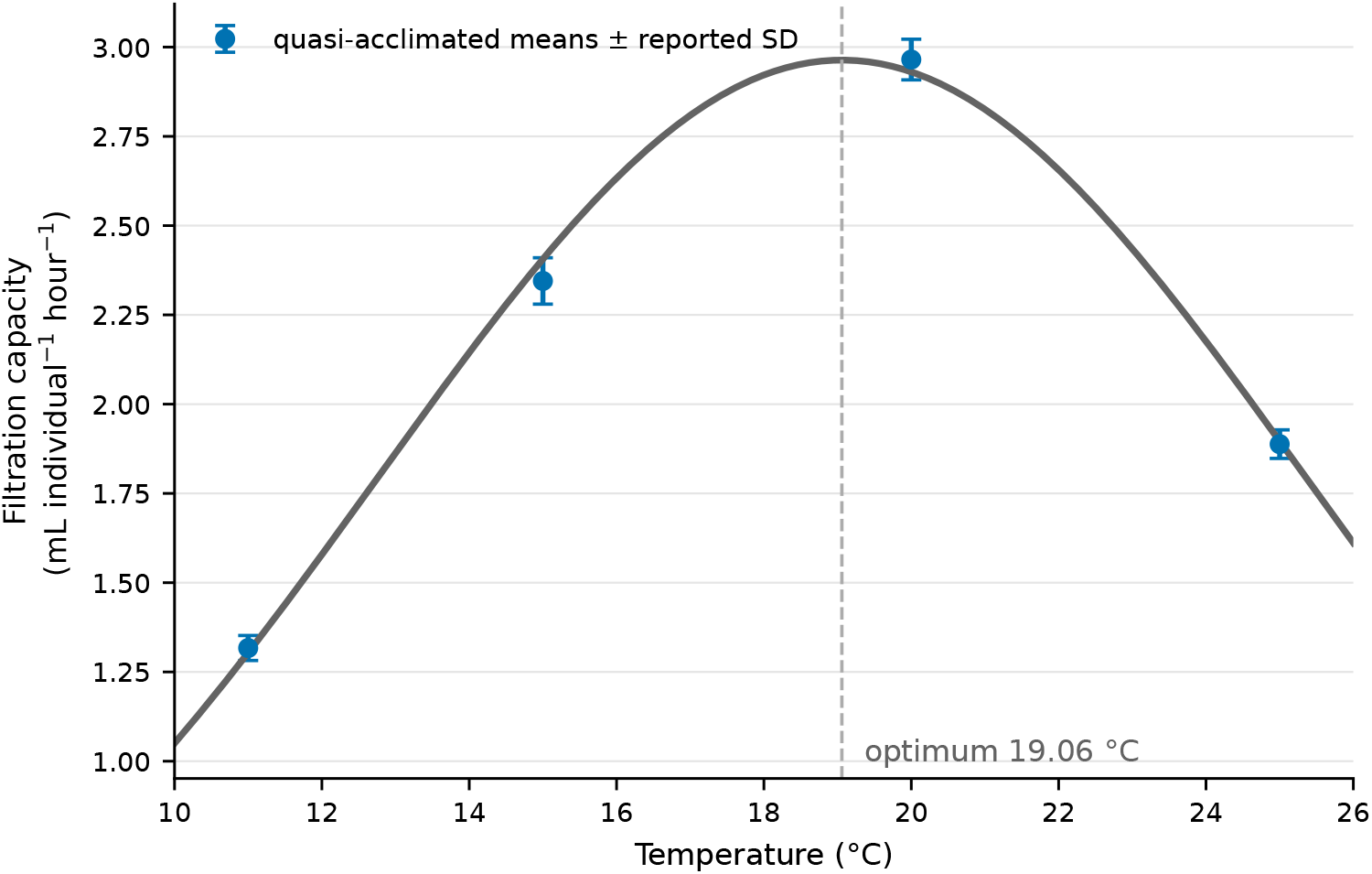
Quasi-acclimated filtration-capacity means and reported standard deviations from the official Müller et al. dataset [21], with the weighted quadratic-exponential fit. The ecological analysis uses only the fitted response relative to its value at 20 °C.

### 5.2 Phase reverses conditional rare-grazer growth

For the primary allocation, both phosphorus ceilings *P*_*i*_*/q*_*i*_ exceed the maximum seasonal carrying capacity 2.55 mg C/L. The producer orbit is therefore light limited throughout the year. Total food biomass ranges from approximately 0.900 to 5.100 mg C/L, while the food factor *h*_*r*_(*t*) changes because the minimum in Eq. (14) switches between food-quantity and phosphorus limitation.

Increasing thermal amplitude broadens the phase response (Figure 2). At *A*_*T*_ = 5 °C, the annual mean ingestion coefficient is 0.68956 day^−1^ and the matched-mean exponent is 0.00840 day^−1^. For comparison, holding temperature at its annual mean gives *α*(20) = 0.8 day^−1^ and an exponent of 0.04979 day^−1^. Replacing this rate by the annual mean 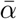 = 0.68956 lowers the phase-independent term by 0.04138 day^−1^ before timing covariance is added. Thus, under this proxy, even a favorable seasonal phase remains below the constant-20 °C result.

**Figure 2:**
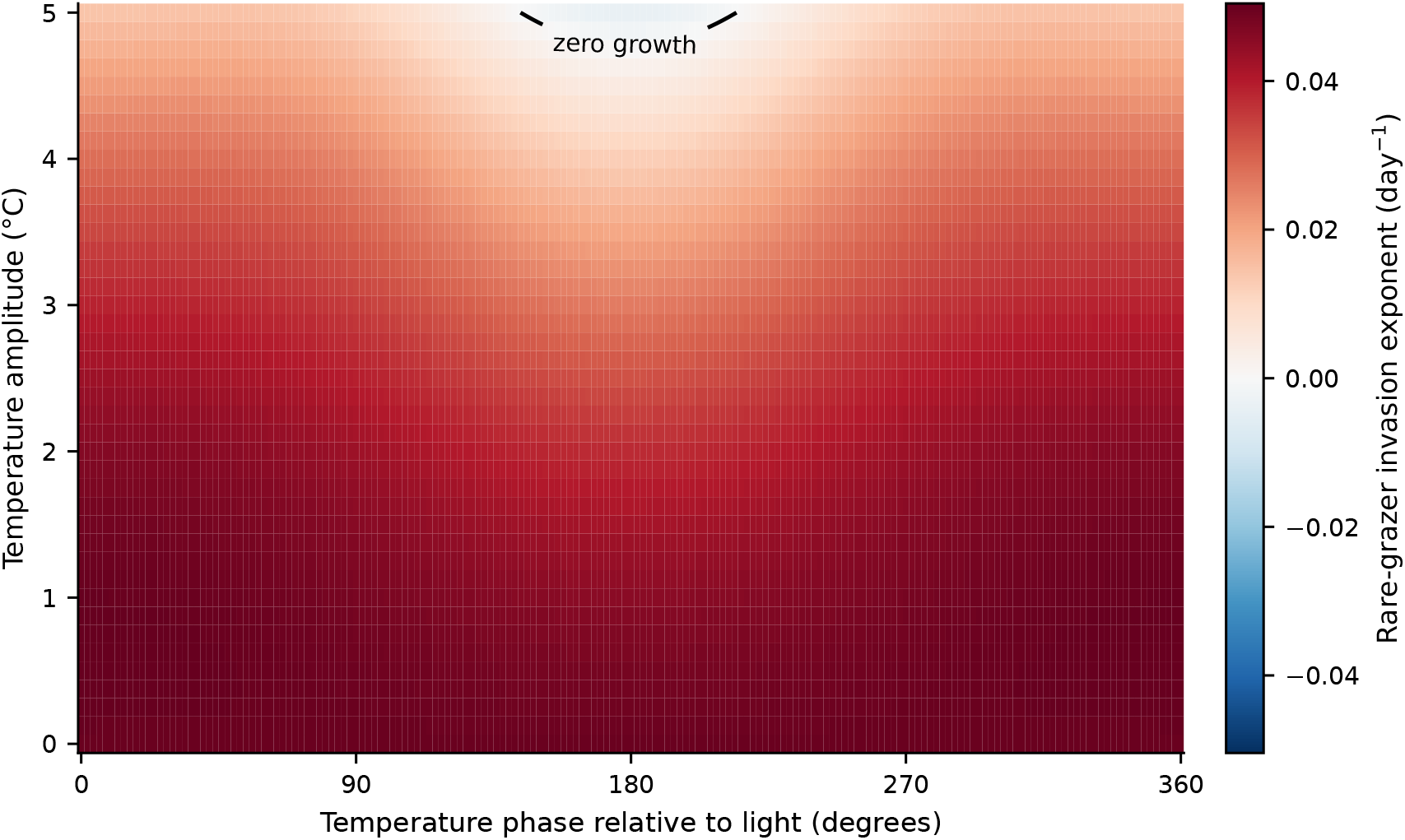
Conditional rare-grazer invasion exponent over temperature amplitude and light– temperature phase for *K*_0_ = 1.5, *ε* = 0.7, and *r* = 0.45. The black contour marks zero annual log growth. Every horizontal row preserves the same marginal temperature distribution across phase.

Nevertheless, phase changes the conditional exponent from approximately −0.00382 day^−1^ near *ϕ* = 179.25° to 0.01486 day^−1^ at two nearly tied favorable phases near 42° and 316°. Their sampled exponents differ by less than the time-quadrature error, so we do not interpret either sampled phase as a unique optimum. The corresponding unfavorable and favorable annual multipliers are 0.248 and 227. Thus, the rare grazer loses about three quarters of its biomass per year in the unfavorable linearized case and increases strongly in the favorable case. These are infinitesimal multipliers, not predictions of unrestricted nonlinear growth.

The covariance decomposition explains the reversal. The constant-mean-ingestion term is independent of phase and is positive in the primary case. The unfavorable phase produces a sufficiently negative covariance between thermal ingestion and the stoichiometric food factor to change the exponent’s sign. Conversely, the favorable phase aligns higher ingestion with more favorable food conditions. Figure 3 shows the resulting difference in cumulative rare-grazer log growth over an otherwise identical annual cycle.

**Figure 3:**
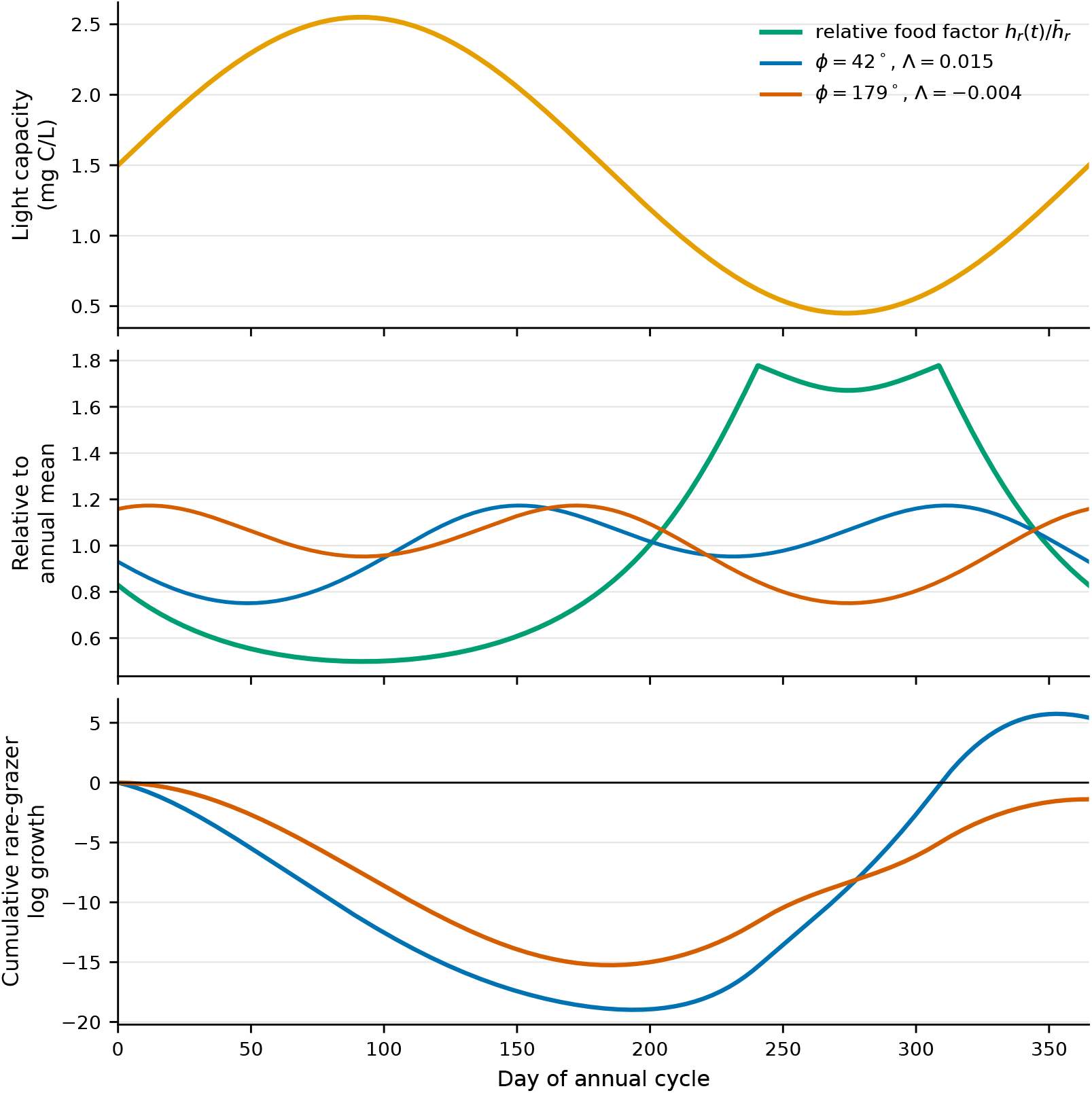
Mechanism for the favorable and unfavorable phases at *A*_*T*_ = 5 °C. Top: annual carrying capacity. Middle: the food factor and temperature-dependent ingestion coefficient, each divided by its annual mean so their timing can be compared on one scale. Bottom: cumulative integral of *α*_*ϕ*_(*t*)*h*_*r*_(*t*) − *d*_*y*_. The endpoint divided by 365 is Λ(*r, ϕ*).

### 5.3 Dependence on the grazer-free phosphorus allocation

Phase sensitivity is not uniform across the nonisolated boundary family (Figure 4). Theorem 3.1 guarantees quota admissibility for all *r ∈* (0, 1) in the numerical light range: *K*_min_ *≥* 0.42 and hence *Q*_*i*_(*t*) *≤* max*{*0.0038, 0.03*/*0.42*} <* 0.07143 *<* 2.5. Every plotted allocation satisfies the numerical quota specialization 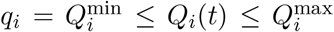; across the allocation and mean-light grids, the quota range is approximately 0.0038–0.0679 mg P/mg C. At *K*_0_ = 1.5, *ε* = 0.7, and *A*_*T*_ = 5 °C, phase changes the sign of Λ for approximately the central allocation range *r* = 0.26–0.74 on the sampled grid. More extreme allocations give positive invasion for every sampled phase. Across all 91 allocation values, 53.8% have both positive and negative phase outcomes. This percentage summarizes a deterministic grid, not a probability distribution over boundary states.

**Figure 4:**
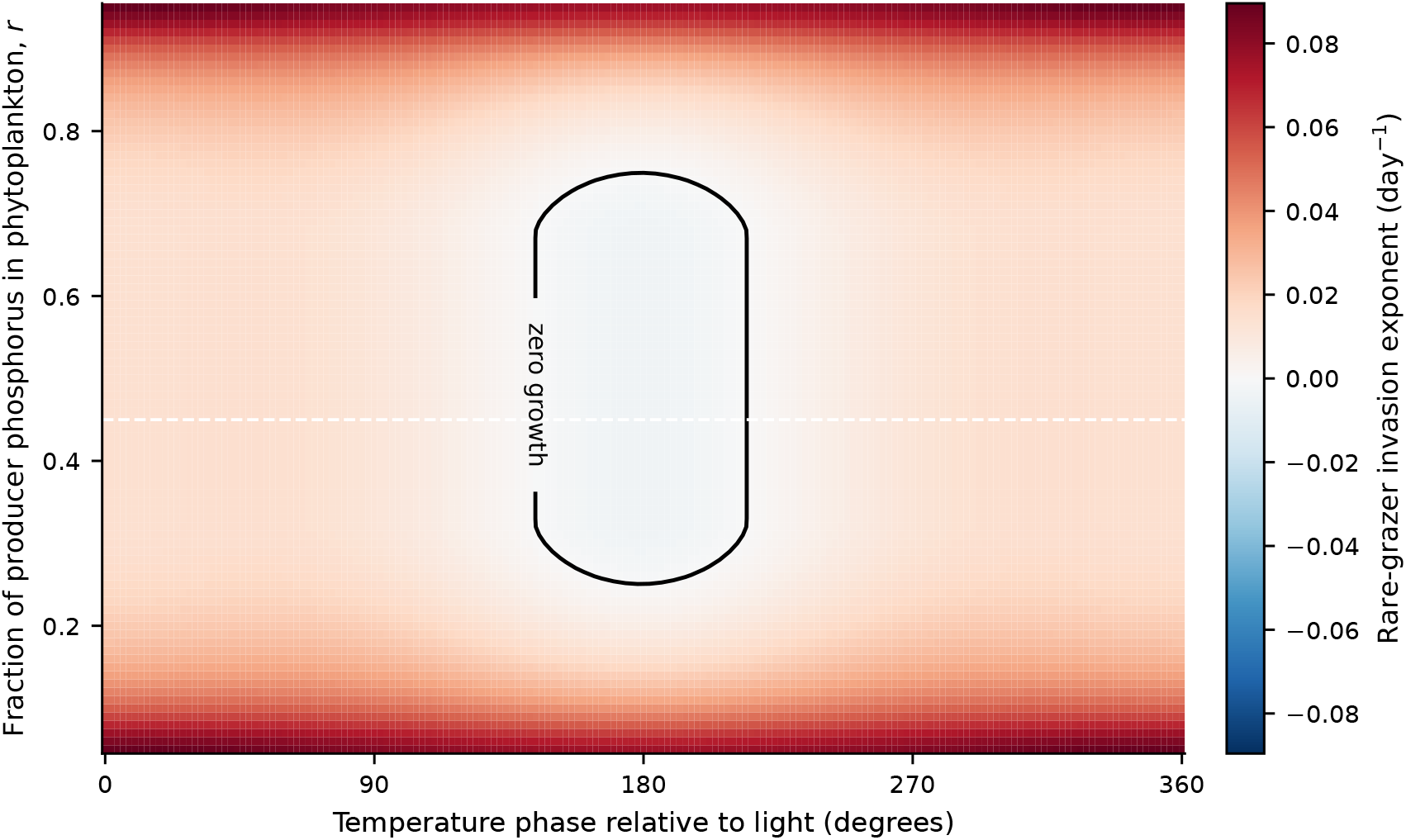
Conditional invasion exponent over phosphorus allocation *r* = *P*_*x*_*/P*_tot_ and phase at *K*_0_ = 1.5, *ε* = 0.7, and *A*_*T*_ = 5 °C. The dashed line marks the primary allocation *r* = 0.45. Different rows are distinct members of the grazer-free boundary family.

The coarser carrying-capacity scan shows that the conditional classification also changes with mean light. At *K*_0_ = 1.4, all sampled phases support invasion over the tested allocations. At *K*_0_ = 1.5 and 1.6, central allocations are phase sensitive. At *K*_0_ = 1.7, a central band has negative exponents for all sampled phases, shoulder allocations remain phase sensitive, and the most asymmetric allocations remain positive. These grid classifications locate regimes for further continuation; their endpoints are not analytical thresholds.

### 5.4 Numerical checks

An independent annual-quadrature check splits the integral at the two food-quality switches and refines the uniform time grid from 0.25 to 0.03125 day at the three reported phase extrema. The original grid differs from the split adaptive integral by less than 7.4 *×* 10^−8^ day^−1^, and the finest grid differs by less than 1.1 *×* 10^−9^ day^−1^. The signs and reported rounding are unchanged. The original quadrature error also explains the residual floor in the full-system rare-grazer comparisons.

The quadrature-initialized primary orbit closes to 1.1 *×* 10^−14^ in maximum absolute biomass. Independent DOP853 and LSODA integrations differ from it by at most 3.4 *×* 10^−11^ and 1.6 *×* 10^−10^, respectively. The covariance identity is satisfied to floating-point precision on every saved grid point. Full-system perturbations at three rare-grazer densities reproduce the positive and negative linear exponents to less than 10^−6^ day^−1^; recovered pelagic-phosphorus minima are at the 10^−16^ numerical scale. These checks support the stated conditional calculation, not global claims about all full-system trajectories.

## 6 Discussion

Within the representative parameter regime, the analysis demonstrates a precise form of seasonal nutritional mismatch. Shifting temperature relative to light changes neither the annual light series nor the annual temperature distribution, yet it reverses rare-grazer annual growth. The covariance identity makes the mechanism explicit: a positive mean-rate control can be overcome when high thermal ingestion occurs during unfavorable stoichiometric food conditions. This agrees with earlier evidence that temperature–food covariance affects ectotherm performance [8] and that storage, frequency, and resource-peak phase affect consumer responses [9]. The contribution here is the conditional boundary calculation in a closed-phosphorus, two-producer habitat model, not the first demonstration of a phase effect.

This result extends rather than repeats the light-seasonality preprint. The earlier study examined population trajectories and bifurcation structure under changes in carrying capacity [19]. Here, the main object is a conditional Floquet multiplier on a newly characterized annual boundary family. The annual reduction clarifies an issue that finite simulations can hide: the preprint’s grazer-free continuum does not collapse to a single resident cycle under forcing. The resident food trajectory can depend on how conserved producer phosphorus is partitioned between habitats when a producer’s phosphorus ceiling constrains growth. Where both producers remain light limited throughout the year, their biomass trajectories and the conditional exponent are independent of allocation, even though their quotas differ. Indexing by allocation captures both regimes.

The result should be interpreted narrowly. Positive conditional invasion does not prove uniform persistence or coexistence, and negative conditional invasion does not prove global extinction. The boundary family has a neutral allocation direction, and the full model may contain other resident trajectories or interior attractors. The large favorable annual multiplier also applies only while the grazer remains rare; density-dependent grazing changes the resident trajectory once grazer biomass is appreciable. Moreover, the favorable 42° trajectory reaches cumulative log growth of approximately −19 within the year, leaving about 6 *×* 10^−9^ of its initial density before recovery. Positive deterministic annual growth therefore does not imply reliable establishment of a finite population through this seasonal bottleneck.

The thermal pathway is intentionally limited. Müller et al. measured filtration capacity, not the maximal ingestion coefficient of this model [20]. Their four quasi-acclimated means constrain a relative unimodal curve only weakly, and the experiment does not identify how producer growth, grazer mortality, assimilation efficiency, or nutrient demand change with temperature. Temperature– stoichiometry experiments and models show that these additional pathways can matter [6, 7, 31, 32]. A recent preprint likewise develops a temperature-dependent stoichiometric herbivore–autotroph model and reports effects of herbivore nutritional traits on persistence and stability [34]; it does not study phase-shifted seasonal forcing or the present two-habitat boundary multiplier. The present calculation therefore identifies a testable timing mechanism under an ingestion-only proxy, not a calibrated climate projection.

Nor do the present comparisons show that separate benthic and pelagic dissolved pools uniquely cause the phase reversal. Closed-nutrient two-producer/one-consumer stoichiometric models already exhibit enrichment-driven equilibria, cycles, and bifurcations [23]. Other components of the present structure also occur separately in prior work: stoichiometric patches [22], coupled pelagic– benthic algae [24], variable-quota aquatic producer competition [26], and phytoplankton–periphyton systems with a shared grazer [25, 27]. McConnell’s recent dissertation treats two dynamic producer quotas and one shared consumer around a common nutrient pool [28]. Recent marine models also combine two phytoplankton resources, one grazer, variable elemental allocation or food-quality-sensitive grazing, and temperature-sensitive or seasonal processes [29, 30]; they do not address the present benthic–pelagic conditional invasion problem. A causal structural comparison requires mass-balanced common-pool and fast-quota reductions with all other processes matched. That comparison, thermal-curve uncertainty, alternative temperature pathways, and variation in forcing frequency are the most important next steps. Nutrient storage can create history dependence under thermal forcing [33], but the present ingestion-only model does not by itself establish thermal memory.

Within the sampled allocation and mean-light grids, the conditional sign reversal recurs often enough to motivate empirical and theoretical follow-up. It generates the hypothesis that identical annual temperature and light distributions can yield positive or negative growth from rarity, depending on their relative timing and the resident phosphorus allocation. Paired measurements of temperature, producer biomass, producer P:C ratios, and grazer feeding would help estimate the seasonal food factor and ingestion response. Controlled phase-shift experiments that preserve the annual temperature and light distributions while measuring growth from rarity would test the predicted reversal.

## Acknowledgments

The author thanks Angela Peace for her guidance on the original dissertation research. The underlying model and dissertation were the author’s work, and the previous light-seasonality preprint was developed from that research. AI tools assisted with identifying the omitted extension, auditing the thermal-data provenance, implementing and checking the new calculations, organizing the literature, and preparing this manuscript. The author is responsible for verifying the analysis and text before dissemination.

## Data and code availability

The accompanying reproducibility archive contains the analysis script, official source dataset with checksum, derived CSV files, figures, and numerical-check summary. The script regenerates the reported analysis and figures. No public repository DOI has yet been assigned.

